# Adaptation of retinal discriminability to natural scenes

**DOI:** 10.1101/2024.09.26.615305

**Authors:** Xuehao Ding, Dongsoo Lee, Joshua B. Melander, Surya Ganguli, Stephen A. Baccus

**Affiliations:** Department of Applied Physics, Stanford University; Neurosciences PhD Program, Stanford University; Department of Neurobiology, Stanford University

## Abstract

Sensory systems discriminate between stimuli to direct behavioral choices, a process governed by two distinct properties - neural sensitivity to specific stimuli, and neural variability that importantly includes correlations between neurons. Two questions that have received extensive investigation and debate are whether visual systems are optimized for natural scenes, and whether correlated neural variability contributes to this optimization. However, the lack of sufficient computational models has made these questions inaccessible in the context of the normal function of the visual system, which is to discriminate between natural stimuli. Here we take a direct approach to analyze discriminability under natural scenes for a population of salamander retinal ganglion cells using a model of the retinal neural code that captures both sensitivity and variability. Using methods of information geometry and generative machine learning, we analyzed the manifolds of natural stimuli and neural responses, finding that discriminability in the ganglion cell population adapts to enhance information transmission about natural scenes, in particular about localized motion. Contrary to previous proposals, correlated noise reduces information transmission and arises simply as a natural consequence of the shared circuitry that generates changing spatiotemporal visual sensitivity. These results address a long-standing debate as to the role of retinal correlations in the encoding of natural stimuli and reveal how the highly nonlinear receptive fields of the retina adapt dynamically to increase information transmission under natural scenes by performing the important ethological function of local motion discrimination.

## 1 Introduction

A central question of sensory neuroscience is how the performance of neural circuits is adapted to the statistics of natural stimuli.^1^ Normative approaches, including efficient coding theory, use theoretical methods to derive how optimal sensory circuits should function.^2–4^ Although such theories have revealed that simple receptive field properties such as linear stimulus sensitivity or thresholds match the expected optimum,^5–8^ they rely on strong and inaccurate assumptions such as Gaussian stimulus statistics and independent neural noise, and have not accounted for complex nonlinear encoding properties.

A second approach has been to measure information transmission under natural and artificial statistics, either in the brain or in behavior.^9–13^ However, such analyses are limited to the stimuli presented in finite experiments, and also give no direct insight into the internal computational properties or mechanisms that generate this increased efficiency.

Here we take a new approach using a mechanistically interpretable network model of the neural code of the retinal ganglion cell population that accurately captures both stimulus sensitivity and stochastic correlations during the presentation of natural scenes. We further model the distribution of natural stimuli using methods from generative machine learning. From these models, we calculate how stimuli can be discriminated given any specific stimulus. This approach not only allows us to assess whether natural stimuli are encoded with greater information, but how the properties of sensory tuning and correlated neural noise adapt to contribute to information transmission for each specific stimulus, thus allowing a direct answer to the long-debated question of the role of noise correlations.^14^

We analyzed the manifolds of natural scenes and ganglion cell responses using information geometry to compute the Fisher information, which accounts for distinguishable stimulus changes as embodied by the “just noticeable difference” that is frequently measured psychophysically in humans. This discriminability limits all behaviors, including subcortical reflexive behaviors and perceptual behaviors distinguishing stimuli in distinct categories. We show that the ganglion cell population adapts dynamically to transmit greater information about the manifold of natural scenes than for random changes in the stimulus. Furthermore, we show that a key property of nonlinear visual processing that contributes to this optimization is the discrimination of local motion. We further find that this optimization arises from the dynamic receptive field structure of the ganglion cell population whereas noise correlations reduce, not enhance, information transmission. These results show that the nonlinear visual sensitivity of the retina adapts dynamically to perform an important behavioral function, to discriminate local motion in natural scenes.

## 2 Results

### 2.1 A stochastic model of the retinal code

Discriminability depends on both sensitivity, defined here as the incremental change in the average population response caused by a change in the stimulus, and the stochasticity, defined here as the joint distribution of the population response when the stimulus is fixed. Estimating sensitivity requires a differentiable model that maps the stimulus space to the representation manifold (Fig. 1a), which is composed of the mean activity. To analyze both sensitivity and stochasticity, we created a stochastic encoding model of the retina that captured both the average response to natural scenes and the correlated noise.

**Fig 1:**
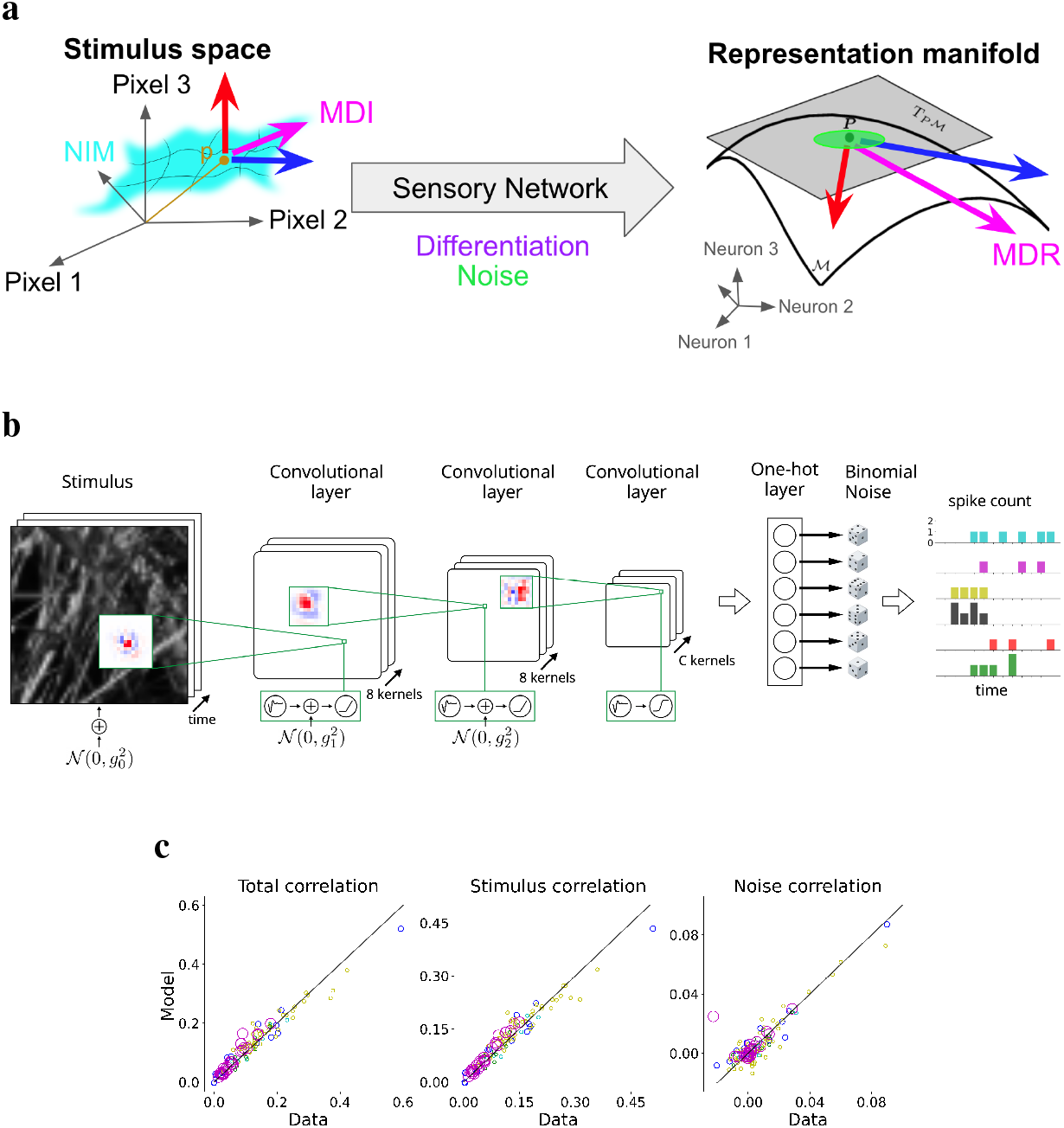
A stochastic model of the retina. (a) Schematic of a general sensory system. With noise in the network, each stimulus corresponds to a cloud of points in the representation space. The stimulus sensitivity for two example stimulus directions (blue and red arrows) and the stochasticity (the green oval) both vary with the specific stimulus. At a particular stimulus point p, the stimulus change that can most easily be discriminated is the most discriminable input (MDI), and is mapped to the most discriminative response (MDR). The tangent plane *T*_*p*_*M* is the local linear approximation to the response manifold obtained by differentiating the network. (b) Architecture of the retinal convolutional neural network model. Layer 1: 8 cell types with 15 × 15 convolutional filters. Layer 2: 8 cell types with 11 × 11 filters. Layer 3: *C* 9 × 9 filters, where *C* is the number of empirically determined cell types. Gaussian white noise is added to the stimulus and the first two convolutional layers, and independent binomial noise is applied after the final nonlinearity. The one-hot layer selects the location of the unit in the final convolutional layer to match that of the recorded neuron. (c) Comparison of total pairwise correlation, stimulus correlation, and noise correlation between neural data and model prediction. Diagonal line is the identity. Each circle corresponds to a pair of recorded neurons in an experimental preparation coded by its color whose radius is proportional to the square root of the test set duration.

We began with a three-layer CNN adopted from Ref.,^15–17^ previously shown to capture the visual sensitivity of the retina under natural scenes and other stimuli (Fig. 1b). Stimuli were images from a natural image database^18^ jittered with the statistics of fixational eye movements. In addition to this model predicting the average natural scene response nearly to within the variability of ganglion cells, the model trained only on natural scenes generalizes to a wide range of ethological computations of motion processing, detection of stimulus patterns and adaptation. Furthermore, the model is mechanistically interpretable in that the activity of internal units is correlated with retinal interneuron responses recorded separately and not used in training. This model was then extended to extrapolate from recorded neurons to entire populations of recorded cell types by adding a third convolutional layer that was then sampled by each of the recorded neurons (see Methods and Ref.^19^).

To assess how the retina discriminates stimuli, it is necessary to have an accurate representation of not only the mean response, but also the variability in single cells and correlations between neurons, of which there are several types. Stimulus correlations arise from common sensory tuning and can be calculated by correlating the activity of cells taken from different trials in response to an identical stimulus. Noise correlations arise from sources other than the stimulus, and are defined as the correlation in the trial-to-trial deviation from the mean response. The total correlation is simply the measured correlation in the activity of two cells, which is also the sum of stimulus and noise correlations. We thus extended the deterministic model of retinal sensitivity to capture the stochastic properties of the ganglion cell population. While freezing all deterministic parameters, stochastic parameters in the model were optimized to fit the single cell variability and pairwise correlations of the recorded ganglion cell population. The only noise sources were independent Gaussian noise added to each cell within the model, with a different noise amplitude for each layer, and binomial noise for each ganglion cell. Simply by adding these few parameters defining independent noise, the full model after optimization successfully captured the mean activity, single-cell variability and correlations between cells including total, stimulus and noise correlations (Fig. 1c, Fig. S1, Fig. S2).

### 2.2 A framework for analyzing stimulus discrimination

A complete knowledge of both the deterministic neural code, or sensory tuning, and a model of the population noise for each stimulus determines the geometry of the manifold of neural responses, as well as the transmitted information and discrimination of stimuli.^20^ Using principles of Information Geometry,^21^ we developed a theory and protocol to analyze the Fisher information of the network in order to determine the discriminability of any changes in the stimulus. For a given stimulus, the network can be considered as a fixed spatiotemporal linear network given that a model consisting only of rectified linear (ReLU) nonlinearities is locally equivalent to an effective linear network,^22^ although this linear approximation changes with each stimulus. The stimulus sensitivity (or tuning) of such a linear network is represented by the Jacobian matrix, the rows of which are the instantaneous receptive fields (IRFs) of each ganglion cell. The IRFs change dynamically over timescales of tens to hundreds of ms reflecting adaptation to the specific stimulus sequence^16^ (Fig. S3). We estimated the noise covariance matrix of the neural response for each stimulus empirically by presenting the stimulus repeatedly to the model. We then calculated the Fisher Information Matrix (FIM) for each stimulus using the Jacobian matrix to capture sensitivity and the covariance matrix to capture stochasticity^21^ (see Methods and Fig. S4).

For each stimulus, we directly computed the stimuli that were most effectively discriminated given the neural code and noise. The most discriminable input (MDI) is the change in the stimulus that the network conveys the most information about in the current state, and was computed as the largest eigenvector of the Fisher Information Matrix.^19^ Additional MDIs could be computed as the 2^nd^ and smaller eigenvectors of the FIM. In addition to the MDI, we computed the response induced by the MDI, which is the most discriminative response (MDR), or the change in the response that would convey the most information to the higher brain about the MDI.

We further examined the relationship between the local computation of MDIs, and discriminability between stimuli separated by larger distances in pixel space (Fig. S5). We found that the ordering of MDIs is preserved across distance, indicating that the local measurement of discriminability is representative for larger stimulus changes. We also computed how different cell types contributed to discriminability, either when considered together or separately (Fig. S6). This analysis showed that different parts of the stimulus contributed to the MDRs for different cell types, indicating that different cell types convey information about salient stimulus features in different regions of the image.

### 2.3 The retina is optimized to encode natural scenes

We then analyzed whether the retina’s ability to discriminate stimuli has been optimized through evolution to the statistics of natural scenes using a generative model of natural images. The set of natural images lies on a manifold with a much smaller dimensionality than the set of all possible images.^1, 23^ For one natural image represented by a point on this manifold, a change in that image along a direction tangent to this manifold represents a natural change of the image, whereas a change with components in an orthogonal direction signifies an unnatural change. We tested whether the retinal code is optimized to discriminate stimulus changes within the natural image manifold.

We parameterized the natural image manifold using a generative adversarial network (GAN) trained on natural images, where the latent code input to the generator serves as a coordinate system of the manifold. This approach has been widely used in image editing research primarily because of the discriminative nature of the GAN’s latent space and the high quality of generated images^24, 25^ (Fig. 2a).

**Fig 2:**
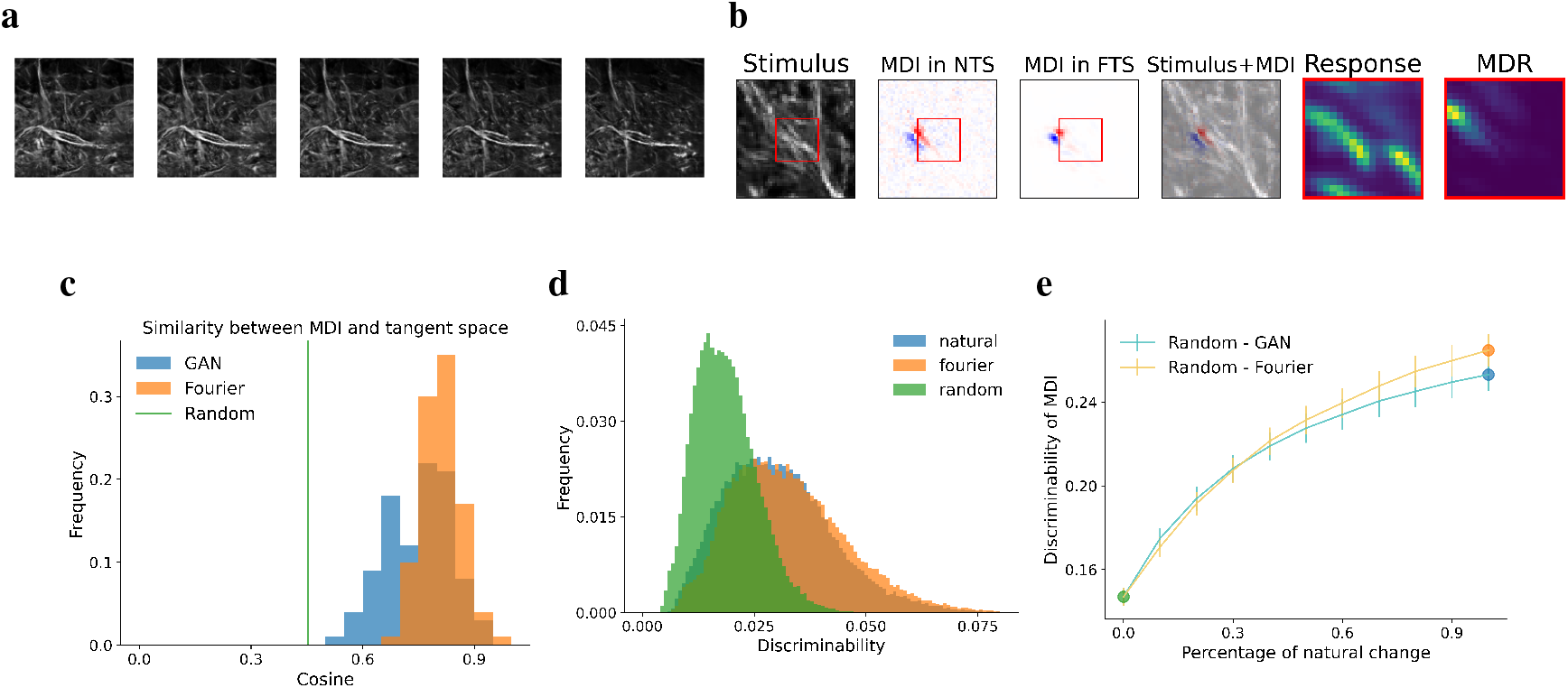
The retina is optimized to encode natural scenes. (a) Example of a sequence of images generated by a GAN by changing the latent code gradually along a straight path. (b) Example of the stimulus alongside the corresponding most discriminable input (MDI). Left to right, Stimulus frame, the MDIs constrained in the natural tangent space (NTS), the full tangent space (FTS), and an overlay of the MDI constrained in the NTS onto the stimulus. Only the first frames of these spatio-temporal vectors are shown given the jittering nature of the stimulus (see the main text). The last two columns show the mean response and the absolute values of the most discriminative response (MDR), averaged over cell-type channels. Red boxes correspond to the unit array in the final layer. (c) Histograms of the cosine similarity between the MDI computed in the full tangent space and its projection onto the natural tangent space, as estimated by both GAN and Fourier analysis across various stimuli. The theoretical value of the expected cosine similarity for a random space (RTS) is shown for comparison. (d) Histograms of discriminabilities of vectors randomly drawn from the NTS (as estimated by both GAN and Fourier analysis) and the RTS, across different stimuli. (e) The discriminability of the MDI constrained to the mixed space spanned by basis randomly drawn from the NTS and the RTS with different ratios.

For each stimulus, we used the GAN to determine the space of natural stimulus changes, known as the natural tangent space (NTS). The NTS is a subspace of the full tangent space (FTS), which is the space of all possible stimulus changes. For each image, we generated stimulus frames in the NTS or the FTS that were then jittered with the same trajectory as the original 400 ms stimulus sequence in order to compute discriminability. We found that over a wide set of stimuli the most discriminable stimulus (MDI) solved in the full tangent space changed little when it was projected into the natural tangent space (Fig. 2b, Fig. 2c). This was remarkable considering that the dimension of the FTS is substantially higher than that of the NTS, indicating that the neural code is optimized to discriminate changes in natural scenes. Examining the structure of the MDIs, we observed many spatially biphasic MDIs split by edges in the stimulus, indicating that discriminability was tuned to local translational motion (Fig. 2b, Fig. S7).

The Fisher Information Matrix can be used to quantify discriminability as a measure of distance between response distributions driven by a change in the stimulus. To compute the discriminability for any stimulus change, we calculated the norm of that vector defined using the metric of the FIM (see Methods), and so compared the discriminability of a large sample of natural and random changes in the stimulus. We found the discriminability of natural stimulus changes was significantly higher than those in a randomly selected space of the same dimensionality (random tangent space, RTS) (Fig. 2d), and that constraining the MDI to mixtures of the natural and random tangent spaces steadily increased discriminability as the space became more similar to the NTS (Fig. 2e). Discriminability under natural scenes also decreased with increasing mean intensity consistent with Weber adaptation,^26^ and also decreased with increasing standard deviation of the stimulus (Fig. S8).

As an alternative simpler method, we instead altered the Fourier spectra of images, as numerous studies have identified the power-law form of Fourier amplitude spectra as a characteristic feature of natural scenes.^1, 27, 28^ An approach often used in psychophysical studies of discriminability is to alter the phases while preserving the power spectrum to either distort a natural image or to blend two natural images naturally.^11, 13^ We approximated natural image changes by fixing the power spectrum of natural images but randomly shifting the phases of Fourier components. This approximation was only applied to changes in the image (the natural tangent space), and the full natural image manifold was used to sample the base stimulus. The results from this approximation were very similar to those using the GAN to model the natural tangent space, confirming that the neural code was optimized to discriminate natural scenes (Fig. 2).

### 2.4 The retina adapts dynamically to discriminate natural stimulus changes

We then examined which properties of the neural code caused the retina to discriminate natural stimuli. We created a simplified version of the CNN model by removing the intermediate nonlinearities while keeping other model parameters and noise levels untouched. This model was the commonly used linear-nonlinear (LN) model, having fixed receptive fields that do not adapt to the stimulus. (Fig. S3). For the LN model, the MDI across the population is stimulus-dependent because different cells are activated by different stimuli. To compare the adaptive properties of stimulus discrimination for the CNN and LN models, we defined for each stimulus the MDI selectivity index as the ratio between the discriminability of the MDI of the current stimulus and the average discriminability for many random natural stimuli. We found that compared to the LN model, the nonlinearities in the retina make the MDI more selective. Therefore, the MDI of the full model changes more dramatically with the stimulus than the LN model (Fig. 3a), indicating that the nonlinear retinal neural code dynamically adapts to natural scenes.

**Fig 3:**
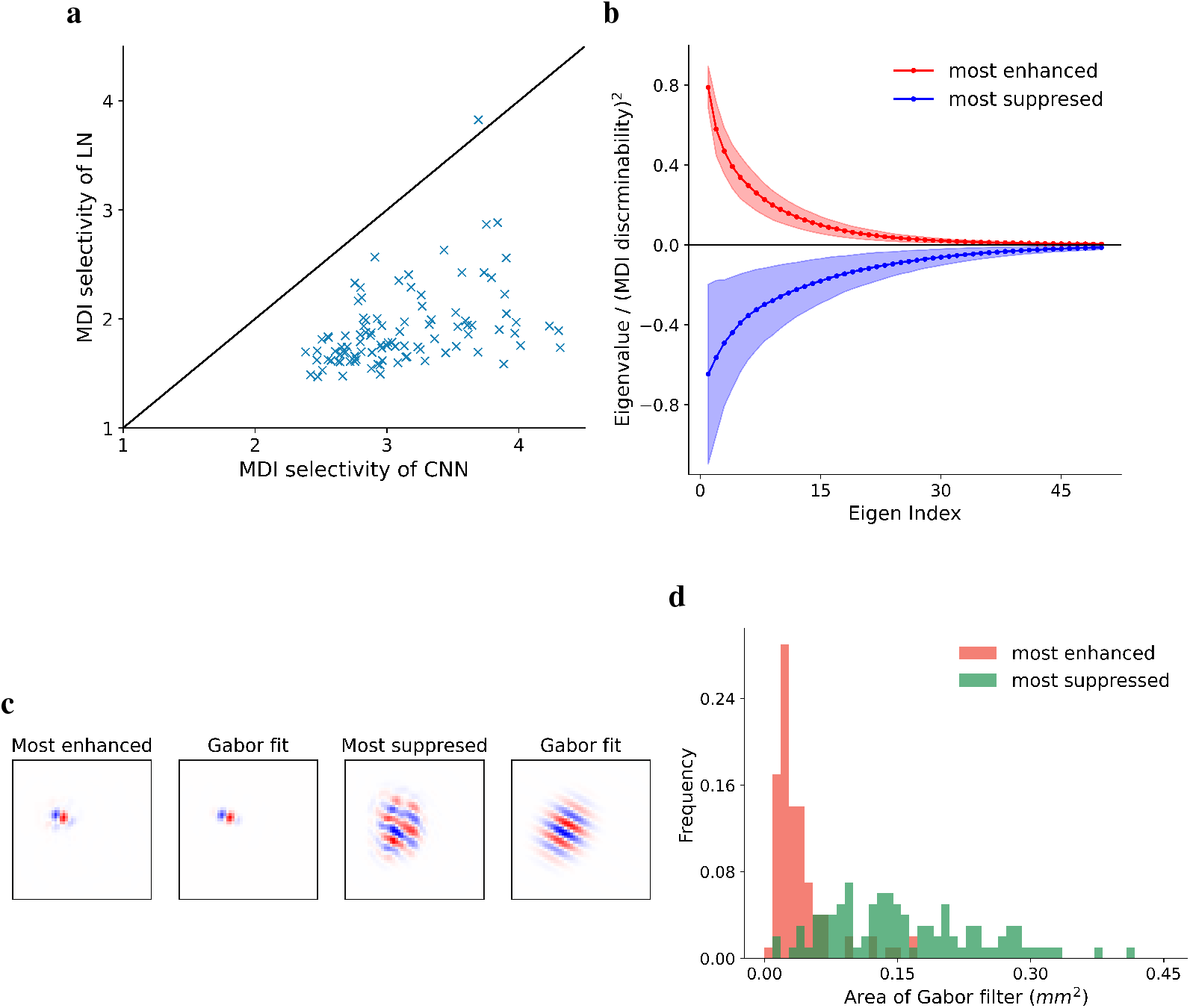
Nonlinearity in the retina enhances the discriminability of local motion but suppresses global motion. (a) MDI selectivity index of the LN model versus the CNN model for different stimuli (see the main text). The black line indicates the identity. (b) Most suppressed and enhanced eigenvalues of the Fisher information difference matrix (FID) normalized by the square of the MDI discriminability over stimuli (mean *±* s.d.) (c) Example of the most positive (enhanced) and the most negative (suppressed) eigenvectors of the FID and the Gabor filters fit to them. (d) Histograms of areas of the Gabor filters fit to the most enhanced and the most suppressed stimulus changes for different stimuli (see Methods).

We then examined which specific stimulus features were enhanced or suppressed by the more complex nonlinearities of the retina. To characterize the differences in how the CNN and LN models discriminated natural stimuli, we defined the Fisher information difference (FID) as the element-wise difference between the Fisher information matrices computed using the full model and the LN counterpart. Eigenvectors of the FID with positive eigenvalues are stimulus changes whose discriminabilities are enhanced by the nonlinearities, while negative eigenvectors are stimulus changes that are suppressed by those nonlinearities (Fig. 3b). We then examined the structure of the eigenvectors of the FID. In the field of computer vision, it has been proposed extensively that motion estimation in natural scenes can be accomplished effectively using the phase of localized sinusoidal functions known as wavelets.^29–32^ As the spatial scale of a wavelet filter increases, the corresponding motion becomes more and more global. We found surprisingly that for most natural stimuli the most enhanced and the most suppressed discriminable stimulus changes can be well approximated by Gabor filters, a type of wavelet (Fig. 3c). The scales of the most suppressed stimulus changes are significantly larger than those of the most enhanced stimulus changes (Fig. 3d), indicating that the nonlinearities in the retina enhance discriminability for local motion but suppress discriminability for global motion. Earlier studies have identified Object Motion Sensitive (OMS) ganglion cells that are sensitive to differential motion as might arise from object motion and insensitive to global motion as might occur from eye movements.^33^ Although these earlier studies used artificial gratings and measured only sensitivity rather than discriminability, our current analysis of the retinal neural code overall points to object motion sensitivity as a key function that drives the dynamic adaptation of discriminability for natural scenes.

### 2.5 Correlated retinal noise limits information

We then tested which properties of the neural code – sensitivity created by dynamic receptive fields, or stochasticity created by correlated noise – contributed to its adaptation for natural scenes by solving for the most discriminable inputs and most discriminative responses when constrained to either natural or random changes in the stimulus. We found that the property of sensitivity was higher for natural than for random changes (Fig. 4a). However, we found that stochasticity was not different when constrained to the natural or random tangent spaces. This finding demonstrated that adaptation of the neural code to natural scenes is driven by adapting receptive fields. This further prompted us to systematically investigate the role of retinal stochasticity, specifically the structure of RGC noise correlations in the processing of information within natural images — a subject of longstanding debate in the field. Multiple scenarios have been advanced where noise correlations are detrimental to information coding.^34, 35^ However, others suggest that noise correlations under certain circumstances may not limit information coding and can even be beneficial.^36–38^ More specifically, theoretical studies have proposed that compared to independent noise, neural correlations driven by noise decrease information when their covariance is aligned with neural correlations driven by signal in the stimulus, whereas noise correlations can increase information when they are structured to be orthogonal to stimulus-driven correlations^14^ (Fig. 4b).

**Fig 4:**
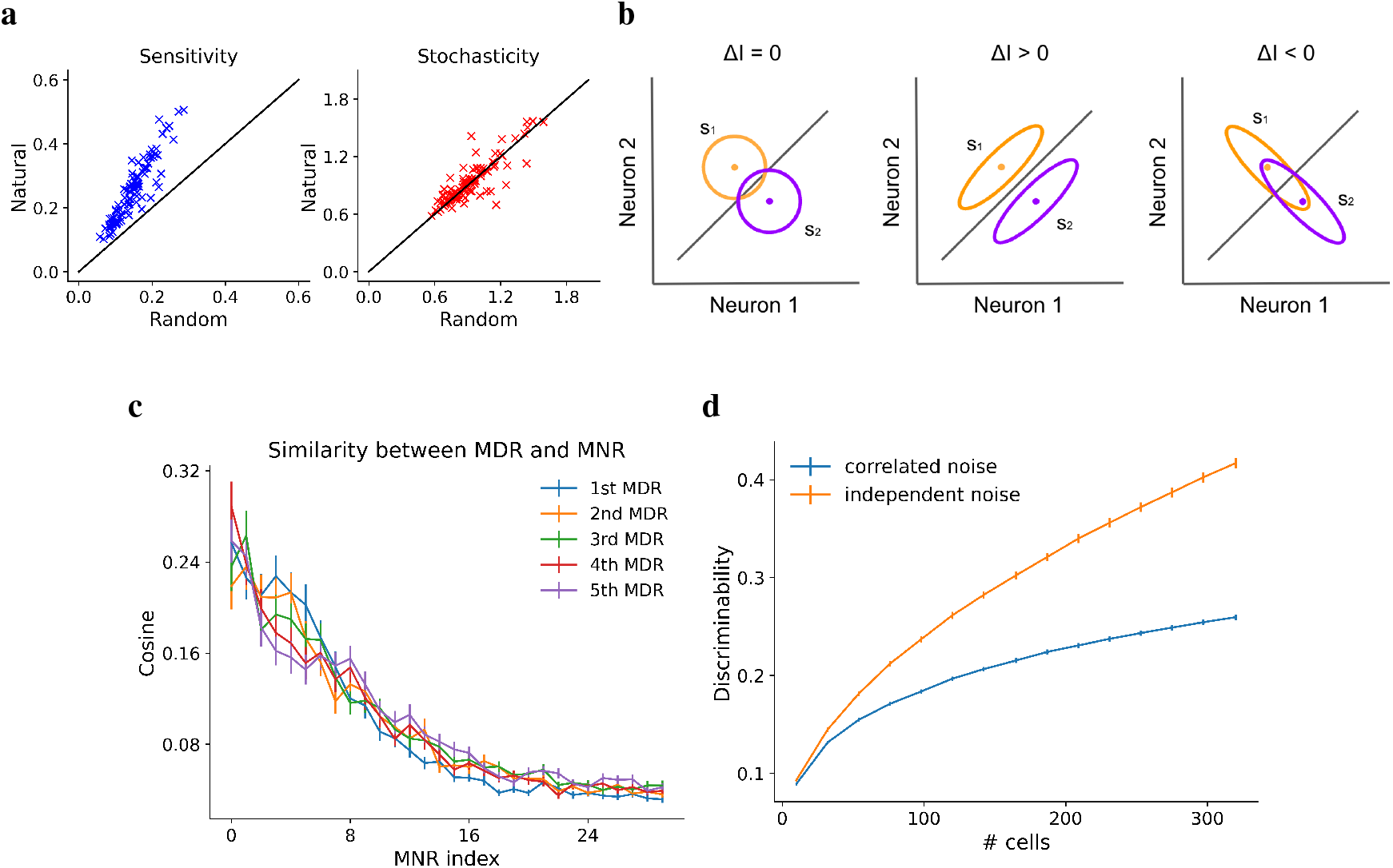
Noise correlations in the retina are detrimental to information coding. (a) Sensitivity and stochasticity of most discriminable inputs constrained to natural and random tangent spaces. (b) Compared to independent noise (left), noise correlations can either increase information (middle) or decrease information (right).^14^ (c) Cosine similarities between the top 5 most discriminative responses (MDRs) and the top 30 most noisy responses (MNRs) averaged over stimuli. (d) Discriminability of MDI for different numbers of output units. The plot for trial-shuffled responses with independent noise is shown for comparison. Output units were randomly selected from the central 500*µm* × 500*µm* region.^39^ MDIs in (c) and (d) were computed in the NTS, similar results are seen for the full tangent space (Fig. S9).

In our model, these two hypotheses could be tested by comparing the most discriminative response (MDR) with the most noisy response (MNR), which was computed as the largest eigenvector of the covariance matrix. If the most discriminative responses are more aligned with the most noisy responses, this implies that noise correlations reduce information transmission. We found that the most discriminative response mode had greater similarity to the most noisy response mode and was orthogonal to modes with low noise (Fig. 4c), implying that noise correlations in the retina are information-limiting correlations. We also performed a more direct test of this conclusion by analyzing trial-shuffled responses, which have independent noise while preserving the single-cell variability. We found that removing noise correlations in this way created responses that were significantly more discriminative than the original model outputs with correlated noise for every population size (Fig. 4d). These observations, in conjunction with our model’s architecture, show that noise correlations do not improve information transmission under natural scenes and may simply arise as a natural consequence of shared circuitry in the retina.

## 3 Discussion

These results show that the receptive fields of ganglion cells dynamically adapt to enhance discriminability to natural stimulus environments, in particular by increasing the discrimination of local motion. The generalizable approach we have created enables an investigation of stimulus discriminability for a broad range of specific stimulus classes and cell types, allowing a direct examination of the relationship between the neural code and information transmission under any set of stimuli. Given the success of machine learning methods at capturing the sensory neural code^16^ and stimulus statistics,^40^ we expect that this approach will reveal similar insight into information transmitted about natural signals in other sensory systems.

Our stochastic model was a simple extension from the deterministic model, created by adding independent noise with no additional effort needed to refit the model’s synaptic weights to capture the correlated noise. We therefore consider the possibility that noise correlations may arise as a natural consequence of independent neural noise and synaptic weights constrained by evolution, rather than as a mechanism to optimize information transmission as previously proposed.^36–38^ Although our analysis indicates that across a large stimulus ensemble and the full ganglion cell population noise correlations limit information, there remains the possibility that noise correlations increase information under some stimulus conditions and for some cell types.

The results in this work are grounded in the examination of infinitesimal stimulus changes. Although this local discrimination ultimately serves to determine discriminability of more distinct stimuli, accurately predicting discrimination across larger stimulus changes will require an analysis of the geodesics under the information metric to navigate between different stimuli.^41^

Visual behaviors of any type are similarly limited by the discriminability of the retina. As additional data is gathered from behavioral and retinal experiments under parallel conditions, a direct analysis of the relationship between information transmission and the neural code will allow an assessment of how the visual properties of different cell types are optimized either alone or in larger populations to discriminate different classes of stimuli in order to support diverse visual behaviors.

## 4 Methods

### 4.1 Retinal recordings and data preparation

A video monitor projected visual stimuli at 30 Hz controlled by Matlab (Mathworks), using Psychophysics Toolbox.^42, 43^ Stimuli had a constant mean intensity of 8.3 *mW/m*^2^. Images were presented in a 50 × 50 grid with a square size of 55 *µm* on the retina. We presented a natural scene stimulus that was a sequence of jittered images sampled from a natural image database.^18^

The responses of tiger salamander retinal ganglion cells from 5 animals were recorded using a 60 channel multielectrode array. Further experimental details are described in Ref.^44^

Spiking responses were binned in 10 ms bins and for the training set were further smoothed using a 10 ms Gaussian filter. Each spatiotemporal stimulus had a duration of 400 ms as we focused on shorter timescales of integration. For each preparation, the training dataset of 60 minutes in total was divided according to a 90%*/*10% train/validation split, and the test dataset consisted of repeated trials to novel stimuli of 60 seconds natural scene.

### 4.2 Details of the stochastic encoding model and optimization

The model (Figure 1b) takes a spatiotemporal visual stimulus as an input and its output is a set of spike counts, one for each neuron in each time bin. Each convolutional filter was implemented by linearly stacking a sequence of 3 × 3 small filters, which outperforms the traditional method in terms of optimization but reduces exactly to a single linear convolutional filter.^45^ A parametric tanh nonlinearity was attached to the last convolutional layer for the purpose of enforcing the refractory period constraint:

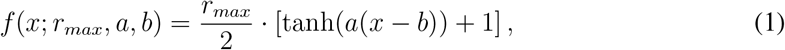

where *a, b* are cell-dependent model parameters, and *r*_*max*_ represents the cell-dependent maximal firing rate.

The essential module that enables us to study retinal population coding is the one-hot layer. Model parameters are fit with the one-hot layer, which is optimized to map a small number of output units of the final convolutional layer to recorded neurons at the correct locations. The one-hot layer is then removed after parameter optimization so that we can analyze the representation of the full population, typically consisting of thousands of units. The rationale behind this method is the mosaic organization of each retinal cell type, in which all ganglion cells of a given type tile the retina with their dendrites.^44^ Formally, the one-hot layer is defined as

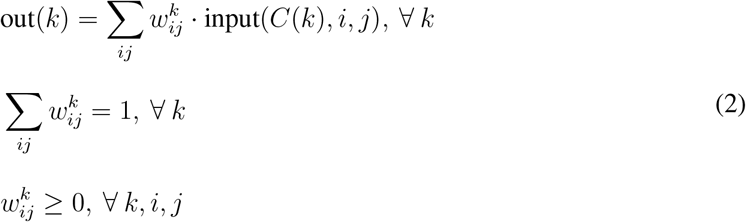

where the input is the output of the last convolutional layer, *k* is the index of recorded neurons, *i, j* (and *p, q* below) are location indices, *w*^*k*^ is the linear combination weight vector, and *C*(*k*) is the neuron-to-channel map. Each *w*^*k*^ will converge to a one-hot array after model training with our one-hot loss function, which is adapted from the semantic loss in Ref.^46^

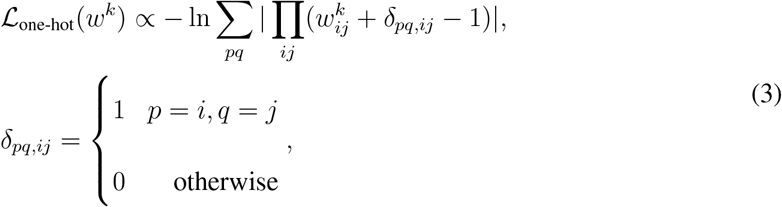

and which reflects the locations of recorded neurons. In order to determine the number of channels in the last convolutional layer and the neuron-to-channel map *C*(*k*), we first fit a model for each preparation assigning a separate channel to each neuron, i.e., *C*(*k*) = *k*. We then computed the cosine similarity between the response vectors of every pair of different channels averaged over stimuli. A standard hierarchical clustering was then applied using the cosine similarity matrix as the affinity matrix. Neurons that were clustered into the same group then shared the same cell-type channel in the final model. The number of clusters as a hyperparameter was explored and the lowest number that did not limit the model performance significantly was chosen.

Gaussian white noise was added to the stimulus to simulate stochasticity in photoreceptors, and was also added to pre-threshold signals in the first two layers to simulate stochasticity in bipolar cells and amacrine cells. We found that Gaussian noise outperforms Poisson noise in these layers, which can presumably be attributed to the noise generation mechanism in the retina and the central limit theorem. The stochasticity of ganglion cell spiking was modeled as a parametric binomial probability mass function (Fig. S2):

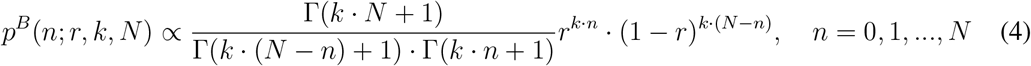

where *n* denotes the spike count in a time bin, *r* is the rate parameter having a one-to-one correspondence with the mean firing rate *µ, k* is the variability parameter to be optimized, and *N* is the cell-dependent upper bound of spike counts in one time bin reflecting the refractory period of that cell, which one can directly determine from data. In practice, for given *k* and *N* we can interpolate the *r* − *µ* mapping and use it to determine the rate parameter producing the desired mean firing rate, i.e., *µ*(*r*) = *E*_*p*(*n*;*r*,*k*,*N*)_(*n*). We stress that the binomial noise on ganglion cells is independent noise, and thus will not qualitatively affect the geometry and the functional implication of noise correlations.

Since the second order statistics are functions of the entire dataset and cannot be decomposed into terms depending on the individual stimulus, we optimized deterministic parameters and stochastic parameters separately. CNN parameters and one-hot parameters {*w*^*k*^} were optimized together with the standard Poisson loss function and ℒ_one-hot_ to fit mean firing rates smoothed using a 10*ms* Gaussian filter. The optimization was performed using ADAM^47^ via Pytorch^48^ on NVIDIA TITAN Xp, GeForce GTX TITAN X, GeForce RTX 3090, TITAN RTX, TITAN V, and GeForce RTX 2080 Ti GPUs. The network was regularized with an L2 weight penalty at each layer and an L1 penalty on the model output. Then tanh parameters of the final nonlinearity were fitted using least squares regression. Then we optimized stochastic parameters while freezing all deterministic parameters. Variability parameters *k* were optimized by maximizing the log-likelihood of recorded spike counts based on empirical firing rates. Standard deviations of Gaussian noise *{g*_0_, *g*_1_, *g*_2_*}* in each layer were optimized through grid searches in order to best fit empirical noise correlations and stimulus correlations by minimizing the smoothed and weighted average of mean squared errors.

### 4.3 Generative adversarial network

The generator of a trained GAN acts as a mapping from latent space to pixel space, denoted as *G* : *Ƶ* → ℐ. Consequently, the differentiation of the generator, *dG*, is a linear transformation between the tangent spaces of these two spaces. We approximated the natural tangent space as the image of *dG* for each generated image.

We follow the same architecture of deep convolutional generative adversarial networks (DC-GAN)^24, 49^ with the image size 1 × 128 × 128. We trained the network using images sourced from the same natural image database that provided the natural stimuli for the retinal recordings.^18^ To prevent mode collapse — a common issue where the network generates a limited variety of outputs — we implemented minibatch discrimination techniques.^50^ We experimented with a range of latent dimensions from 100 to 512, finding that our results were consistent regardless of the specific dimensionality chosen.

### 4.4 Theory of Information Geometry

Let ***x*** denote the stimulus vector and ***y*** denote the spike count vector. Each stimulus induces a conditional probability distribution of sensory neural responses *P* (***y***|***x***) and mean firing rates ***µ***(***x***) := *E*[***y***|***x***]. The representation manifold is defined as the manifold of mean neural responses induced by stimuli: {***µ***(***x***)}. In our case, ***µ***(***x***) is modeled by the noiseless CNN and *P* (***y***|***x***) is modeled by the full stochastic model.

We here define several important matrices. The sensitivity structure is characterized by the Jacobian matrix 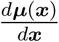. The stochasticity structure is characterized by the noise covariance matrix **Σ**(***x***) := *E*[(***y***−***µ***(***x***))***·***(***y***−***µ***(***x***))^*T*^ |***x***]. Assuming Gaussian noise covariance, the discriminability structure is characterized by the linear Fisher information matrix

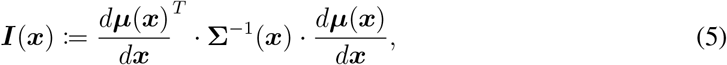

which as an information-geometric Riemannian metric measures the Kullback–Leibler divergence between response distributions induced by infinitesimally close stimuli.^21^ Consequently, the length of a geodesic connecting two stimuli under this information metric measures the discriminability between them, when represented by the neural population.

The rationale for using linear Fisher information rather than the full metric is that (1) this is a good approximation for our model because a neural network consisting only of ReLU nonlinearities is locally equivalent to an effective linear network^22^ and the covariance matrix of a linear network is a constant matrix; and (2) linear Fisher information bounds the performance of unbiased linear decoders which may be more biologically plausible than nonlinear decoders.^51–53^

For any vector ***ξ*** in the stimulus tangent space of stimulus ***x*** such that ||***ξ***|| = 1, the sensitivity and discriminability of it are given by 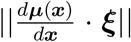 and 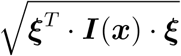, respectively. For any response mode ***a*** such that ||***a***|| = 1 with stimulus ***x***, its sensitivity and stochasticity are given by 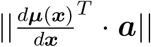 and 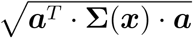, respectively. Furthermore, the MDI and MNR can be obtained as the principal eigenvectors of ***I***(***x***) and **Σ**(***x***), respectively. The MDR is defined as the pushforward of the MDI by ***µ***(***x***): 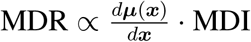.

One can also evaluate the Fisher information matrix and MDI constrained in a subspace of the full stimulus tangent space by

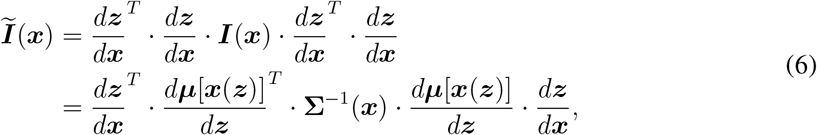

where ***z*** denotes the coordinate with respect to an orthonormal basis of the subspace. Since 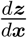 is a semi-orthogonal matrix, eigendecomposing ***Ĩ*** (***x***) simplifies to eigendecomposing 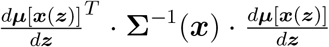, which is more feasible computationally due to the lower dimensionality of the subspace. This method facilitated the computation of MDIs constrained in subspaces discussed in the main text. Additionally, even for computing MDIs in the spatio-temporal full stimulus tangent space, this method was also employed as we proved elsewhere that all MDIs must reside in the subspace spanned by instantaneous receptive fields, i.e., span 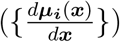.^19^

### 4.5 Statistical measures

Let the neural response be represented by the tensor *R*_*tsc*_ ∈ N denoting the spike count of the *c*-th cell during the *s*-th time bin in the *t*-th trial. Statistical measures used in the main text are defined as follows.

Total pairwise correlation:

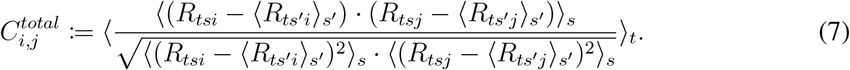

Stimulus correlation:

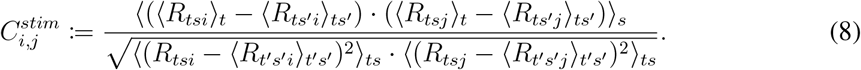

Noise correlation:

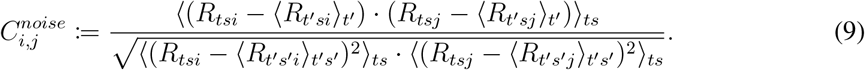

Trial-to-trial correlation:

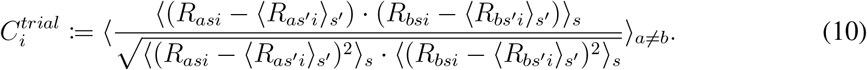

Fano factor:

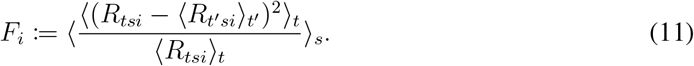

In some previous work, the stimulus correlation is alternatively defined as the average correlation between the neural responses of two cells in different trials. Correspondingly, the noise correlation is defined as the difference between the total correlation and the stimulus correlation. In practice, this set of definitions always produced very similar values to our definitions.

### 4.6 Gabor filter

The Gabor filters we used take the following form:

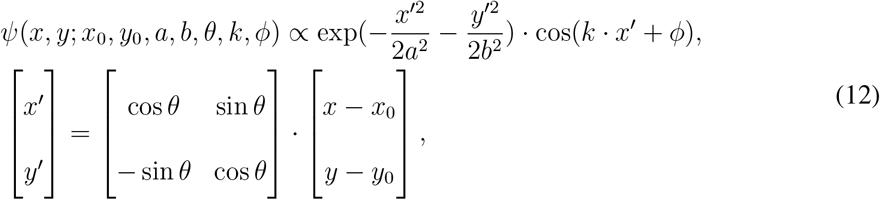

where {*x*_0_, *y*_0_, *a, b, θ, k, ϕ*} is the set of parameters. We optimized these parameters to maximize the cosine similarity. The area of each Gabor filter is defined as the area of the ellipse contour: *πab*.

## Acknowledgments

This work was supported by grants from the NEI, R01EY022933, R01EY025087 and P30EY026877 (SAB).

## Notes

### Competing Interest Statement

The authors have declared no competing interest.

### Summary of Updates

Fig. 3b revised; Supplemental figures reordered; Language polished.

