## supplemental material for "Adaptation of retinal discriminability to natural scenes"

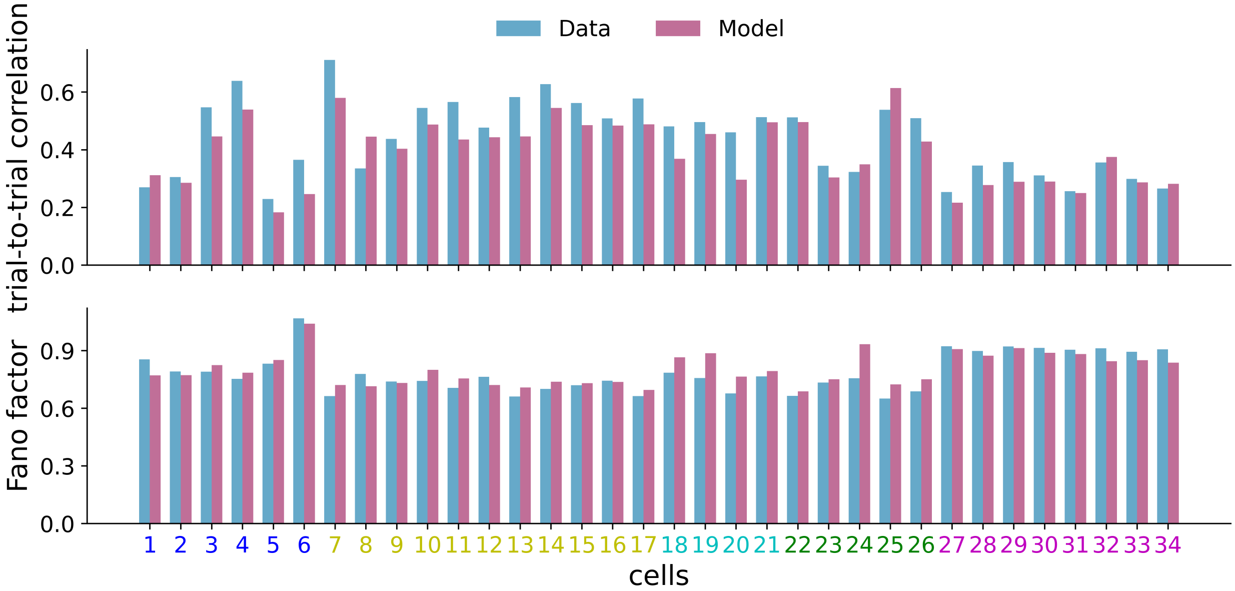


Fig S1: Comparison of single-cell variability measures including the Fano factor and trial-to-trial correlation between neural data and model prediction. Neuron indices share the same color code with Fig. 1c.

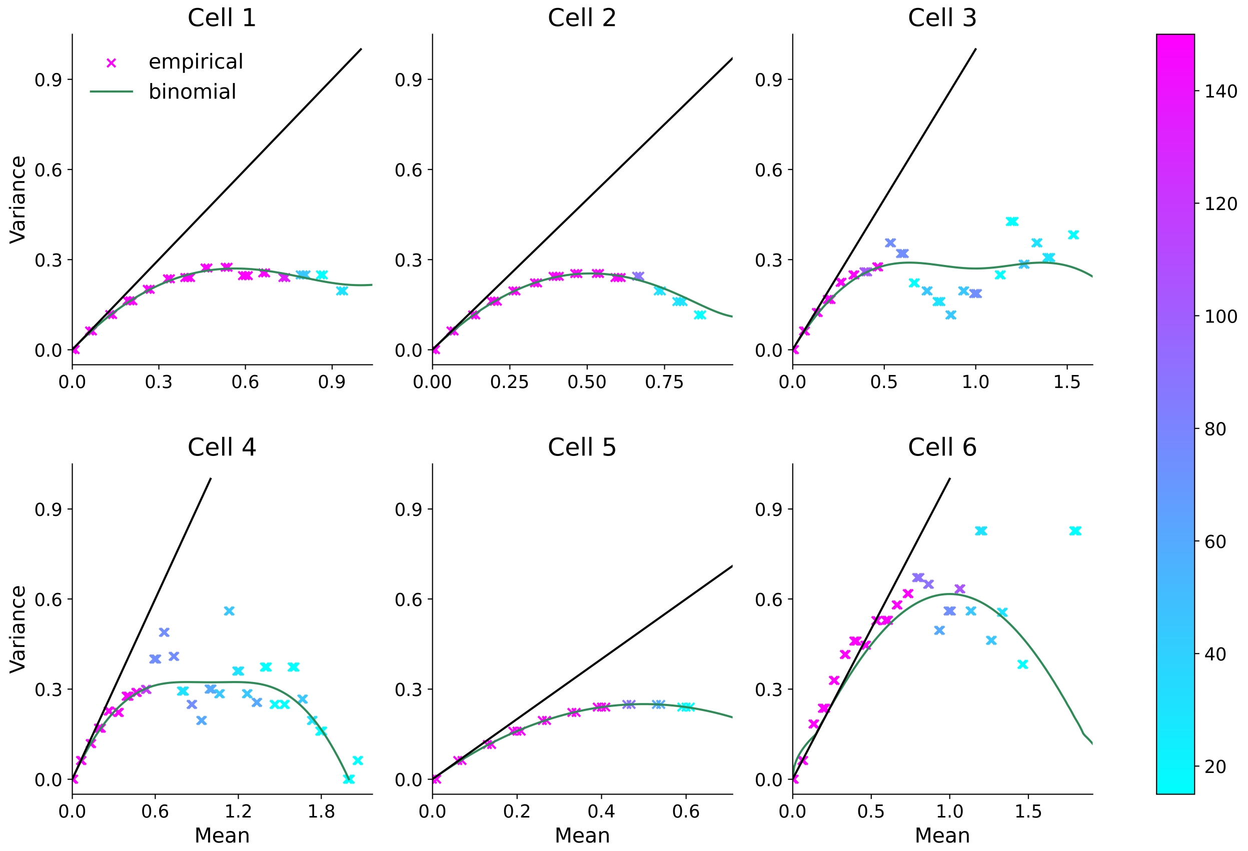


Fig S2: Variance-mean plots for empirical spike counts and the binomial probability mass function after optimization. Each empirical point represents the statistics of samples in all time bins with similar firing rates and its color codes the sample size. Black line is the identity line. Binomial noise captures the empirical relation very well except for some outliers that fluctuate more because of their small sample sizes.

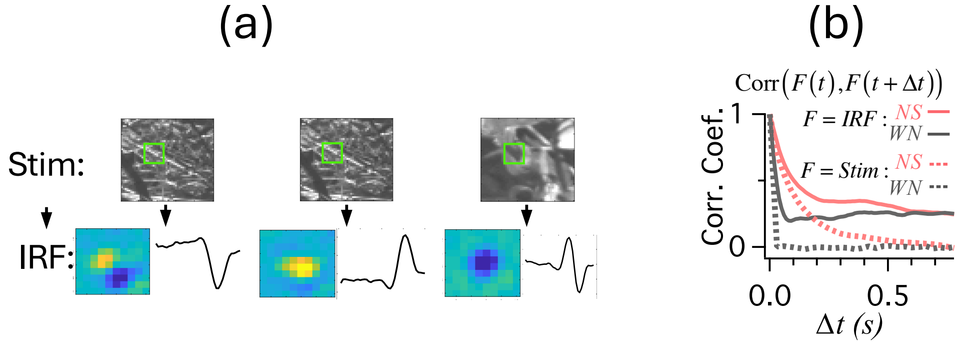


Fig S3: Instantaneous receptive fields change dynamically. (a) For a single ganglion cell, several examples of IRFs computed as the gradient of the model when different natural stimuli were presented. Top shows stimulus frame, bottom shows the spatial and temporal average of the IRF. (b) Average correlation coefficient between IRFs at different times separated by a time interval for natural scenes (NS) and white noise (WN). Also shown for comparison are average correlations between stimulus frames.

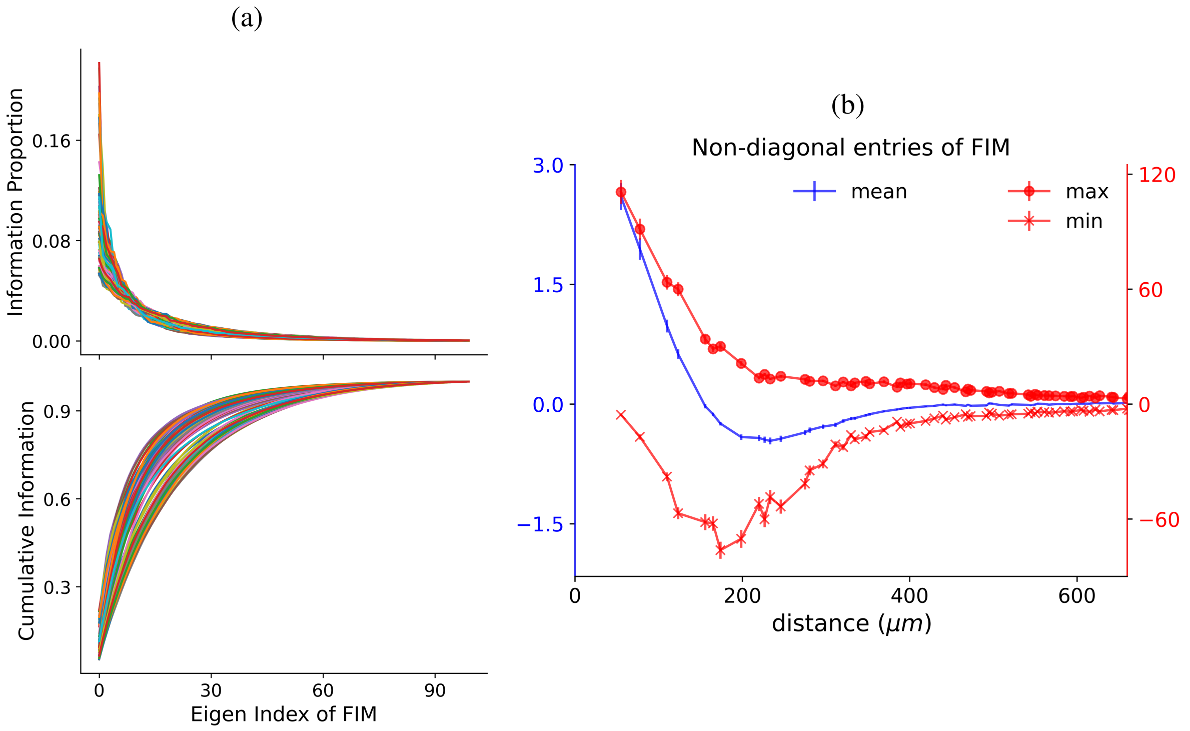


Fig S4: Structure of the Fisher information matrix (FIM). (a) The eigenvalues of FIM ($\lambda_i$) in descending order. Top, the proportion of information carried by each eigenvector: $\frac{\lambda_i}{\sum_k \lambda_k}$. Bottom, the cumulative information proportion carried by the top $i$ eigenvectors: $\frac{\sum_{k=1}^{i} \lambda_k}{\sum_k \lambda_k}$. Each trace corresponds to a distinct stimulus. (b) The blue curve plots the mean value of non-diagonal entries of FIM constrained in the full tangent space (FTS) with the same distance between locations associated to their row and column indices for each stimulus, whereas the red curves plot the maximal and the minimal values. All values are averaged across stimuli.

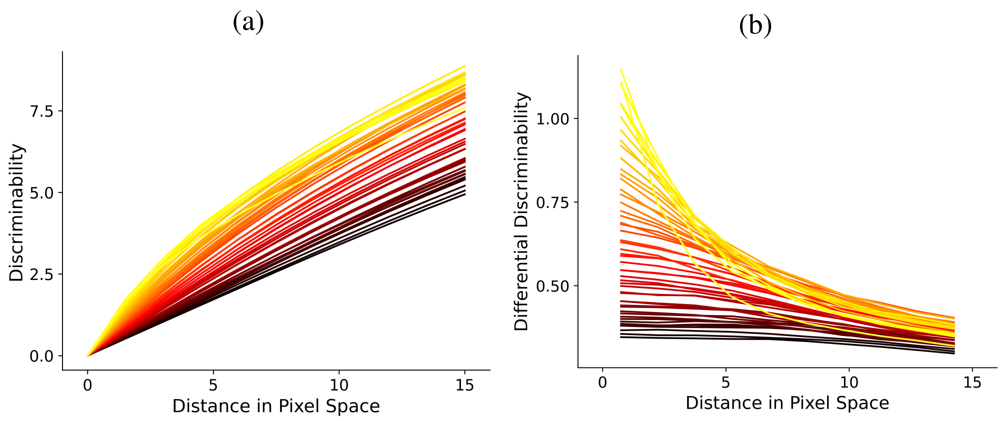


Fig S5: Analysis of nonlocal stimulus changes. (a) Discriminability between the base stimulus and finitely distanced stimuli in the directions of MDIs estimated by integration along straight lines under the information metric. The color codes the eigen index of the FIM at the base stimulus. Values of discriminability are averaged across different base stimuli. (b) Differential discriminability, which is the slope of the curve in (a).

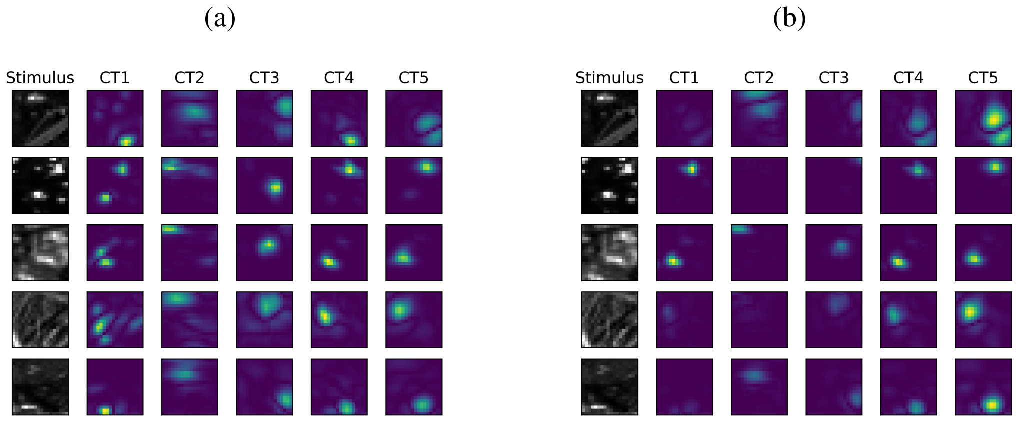


Fig S6: (a) Examples of the most discriminative response (MDR) computed with the presence of distinct cell types for different stimuli. Each column corresponds to a different cell type. (left panel, the central $18 \times 18$ array is shown). (b) Examples of the MDR with the presence of the full population decomposed into cell-type components for different stimuli.

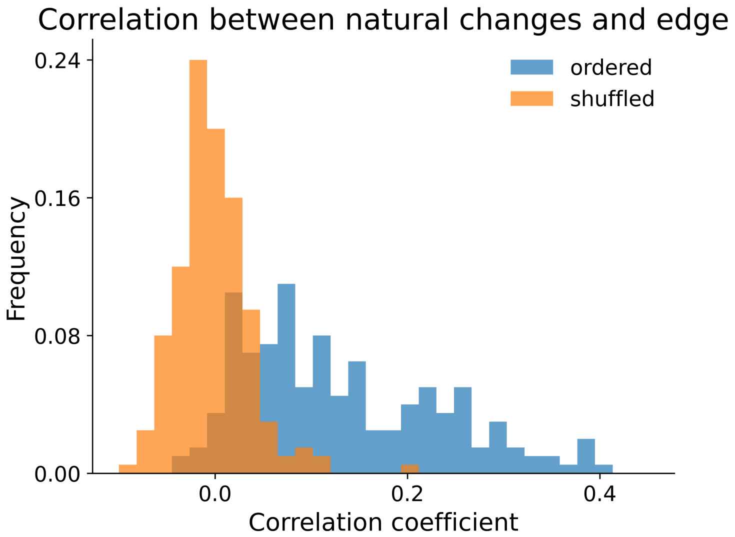


Fig S7: Natural changes produces by a GAN correlate with edges. Histogram of the Pearson correlation coefficient between the absolute values of randomly selected natural changes produced by the GAN and stimulus edges as detected by the Canny edge detector. The histogram for mismatched pairs of stimulus and tangent vector is shown for comparison.

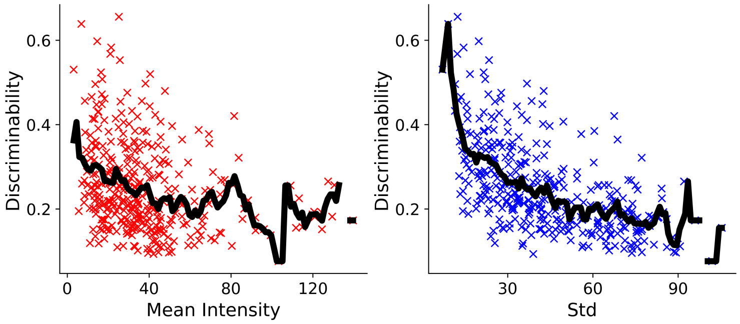


Fig S8: Adaptation of discriminability to mean intensity and standard deviation. The left panel shows the relationship between the discriminability of MDI and the mean intensity of the stimulus image. Each point corresponds to a stimulus. Mean values were plotted with thick black lines. The right panel is similar but for the standard deviation of the stimulus image.

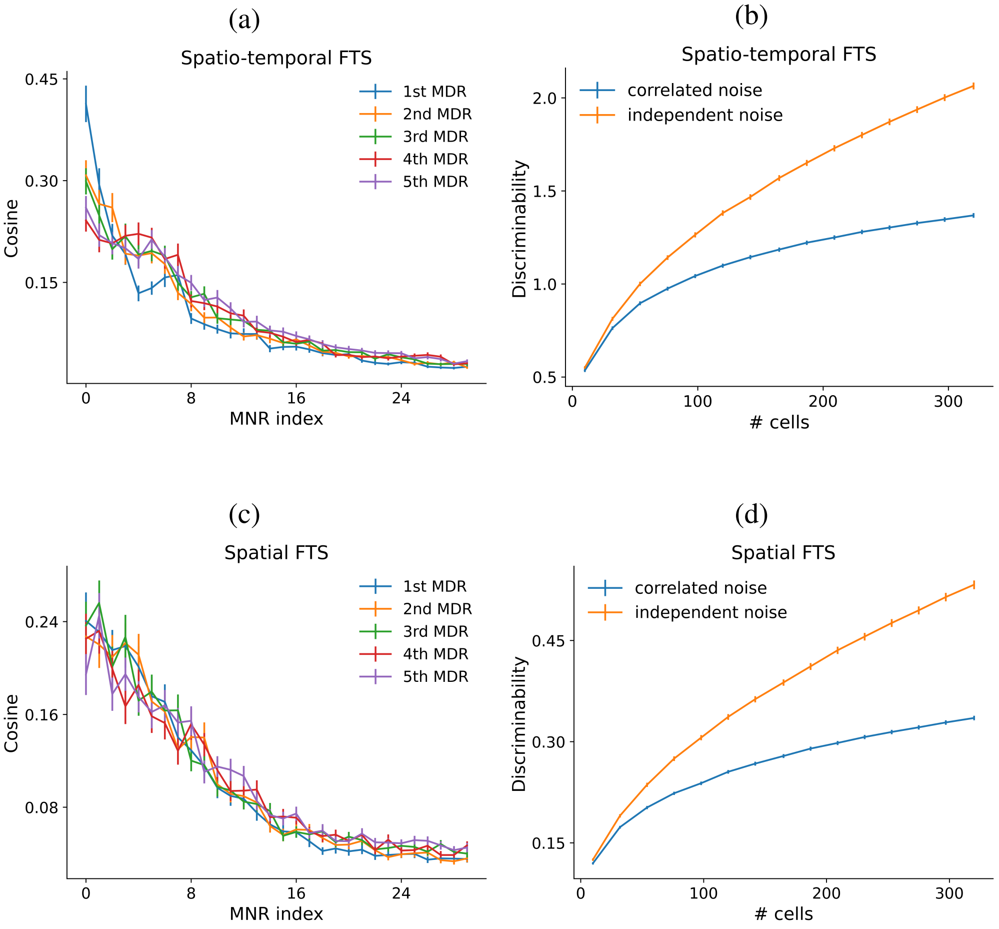


Fig S9: Same plots as Fig. 4c and Fig. 4d but for MDIs constrained to the full tangent space. (a) and (b) are for the tangent space of the full spatio-temporal stimulus space. (c) and (d) are for the tangent space of the full pixel space with the same jittering trajectory as the base stimulus.
